# L-Asparaginase treatment induces reversible immunoregulatory and immunosuppressive effects in non-malignant B cells in a model of T-cell dependent B cell activation

**DOI:** 10.1101/2024.11.27.625617

**Authors:** Amar Hadzic, María Alejandra García-Márquez, Pol Bannasch, Nina Haindl, Katharina Frey, Martin Kirmaier, Maximilian Funk, Anneli Tischmacher, Jingke Tu, Adriano Carboniero, Ludovica Vona, Sabine Oganesian, Hans Schlößer, Werner Schmitz, Michael von Bergwelt, David M. Cordas dos Santos, Sebastian Theurich

**Affiliations:** Cancer and Immunometabolism Research Group, Gene Center, Ludwig-Maximilians-Universität München, Munich, Germany; Department of Medicine III, LMU University Hospital, Ludwig-Maximilians-Universität München, Munich, Germany; German Cancer Consortium (DKTK), Munich Site, and German Cancer Research Center, Heidelberg, Germany; Comprehensive Cancer Center Munich, LMU University Munich, Munich, Germany; Bavarian Cancer Research Center (BZKF), LMU University Hospital, Munich, Germany; Center for Molecular Medicine Cologne (CMMC), University of Cologne, Medical Faculty and University Hospital Cologne, Cologne, Germany; Department of General, Visceral, Cancer and Transplantation Surgery, University of Cologne, Medical Faculty and University Hospital Cologne, Cologne, Germany; Theodor Boveri Institute, Biocenter, University of Wuerzburg, Wuerzburg, Germany

## Abstract

Metabolic reprogramming is critical for immune cell adaptation upon activation to exert full functionality with amino acids being a key metabolic factor. While L-asparagine is a non-essential amino acid it turns out to be conditionally essential in malignant B cells due to defective asparagine synthetase making L-asparaginase a commonly used chemotherapeutic agent. Off-target enzymatic activity including glutaminolysis impacting crucial immune function. However, its effects on healthy B cells remain unclear, therefore in this study, we explored how L-asparaginase modulates the biology and function of CD40-activated B cells, using an in-vitro model. B cells from healthy donors were treated with increasing L-asparaginase concentrations and analyzed for proliferation, immune phenotype, and metabolic changes. Results showed L-asparaginase reduced B cell proliferation and homotypic clustering without inducing apoptosis, instead impairing metabolic pathways, lowering glycolysis and oxidative phosphorylation, and reducing surface markers associated with antigen-presenting cell (APC) function. Functional assays confirmed that L-asparaginase-treated B cells had diminished ability to activate T cells. Supplementing with asparagine or glutamine restored B cell proliferation and function, with glutamine slightly more effective than asparagine. Interestingly, L-asparaginase induced a regulatory B cell phenotype, marked by CD24+CD38+CD27+ expression and increased interleukin-10 and TGF-beta, suggesting a potential immunosuppressive mechanism. These findings indicate that L-asparaginase not only affects malignant cells but also impacts the function of non-malignant B cells, proposing potential therapeutic applications in B cell-driven autoimmune disorders. Further studies are needed to explore its effects at lower, clinically relevant concentrations.

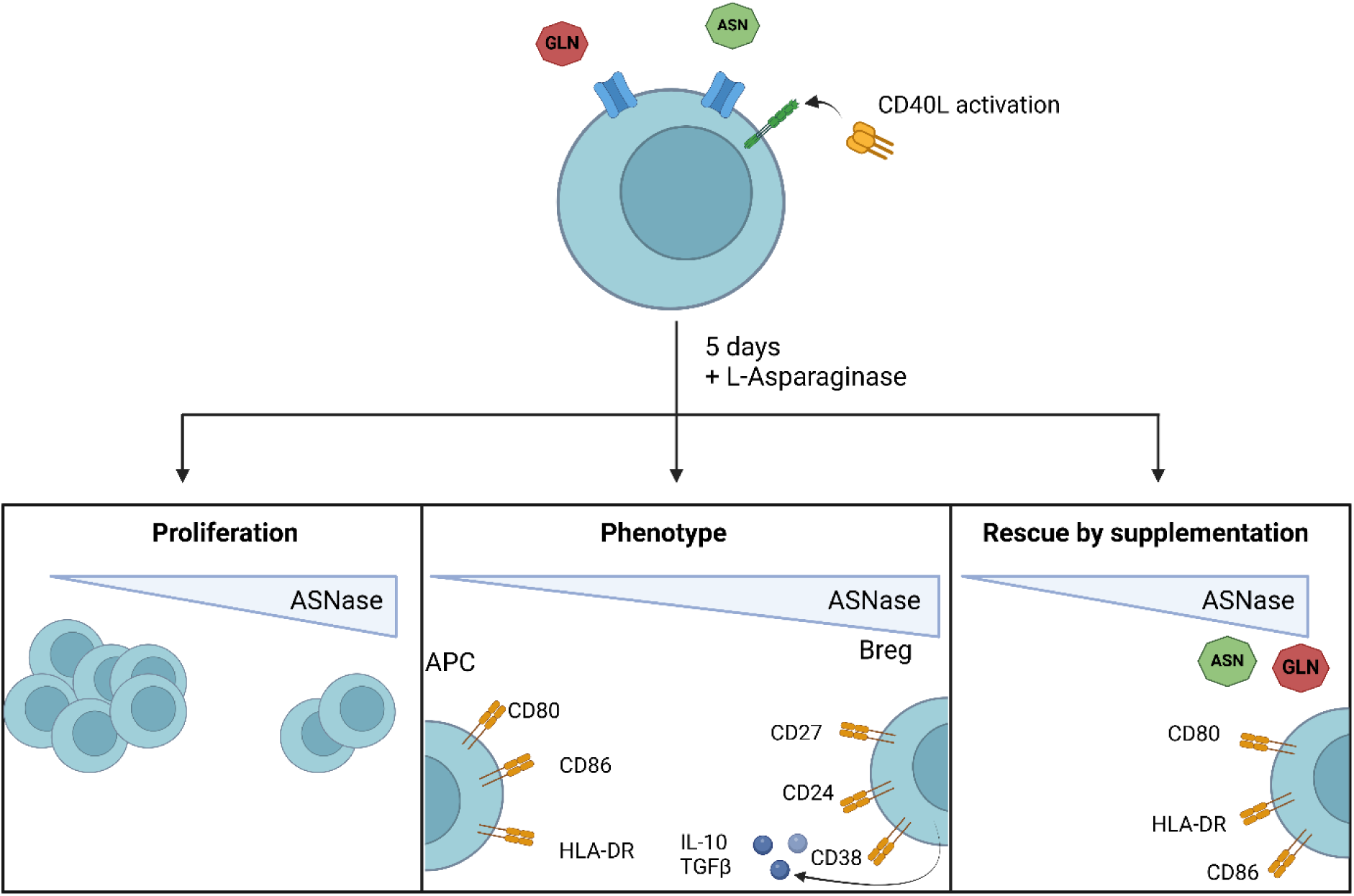

## Introduction

Cellular metabolism plays a critical role in determining immune cell phenotype and function. While metabolic reprogramming is a well-established hallmark of T cell activation it is also observed in B cells, particularly upon activation. In order to meet the increased demands associated with proliferation and effective humoral responses B cell metabolism is upregulated (1). Naïve B cells remain in a metabolically quiescent state, but upon engagement of CD40 on B cells with CD40L on T cells, a cascade of activation is triggered, leading to survival, proliferation, and maturation (2). Thus, leveraging specific metabolic vulnerabilities has proven to be an effective strategy in targeting malignant B cells. Asparagine has emerged as a key metabolic target in the treatment of B cell malignancies due to their lack of functioning asparagine synthetase and reliance on extracellular supply.

In the setting of acute lymphoblastic leukemia (ALL) and lymphoblastic lymphoma (LBL), asparaginase (ASNase) is incorporated into multi-agent treatment regimens to exploit the asparagine dependence of malignant B cells (3, 4). ASNase formulations, including the native form derived from *Escherichia coli* and PEGylated ASNase with reduced immunogenicity and improved pharmacokinetics, deplete asparagine in the serum, leading to apoptosis in asparagine-dependent malignant B cells (5–8). Although the effects of ASNase on malignant B cells are well-documented, its impact on reactive, non-malignant B cells remains elusive. Very few early studies, dating back 60 years, hinted at an immunosuppressive effect for ASNase. Using allograft skin transplant models and murine antibody responses to sheep red blood cells, researchers observed extended graft survival and reduced lymphocyte migration and proliferation in response to ASNase treatment (9, 10). Additionally, B cell function appeared impaired, as indicated by a diminished humoral immune response (11). Building on these early insights, we investigated the effects of ASNase on healthy B cells and the effects on cellular metabolism and phenotype.

Functionally distinct subpopulation form throughout B cell development and maturation. While B cells can perform antigen-presenting function they can also form into regulatory B cells (Bregs), a functionally analogous population to regulatory T cells (Tregs) that serves to modulate immune responses and maintain tolerance. While the Breg subset was initially defined by IL-10-secreting B10 cells, additional markers and cytokines, such as TGF-β, have since been identified, expanding the Breg profile (12–16). Bregs are essential in controlling autoimmunity and hypersensitivity reactions, and their dysfunction can negatively impact Treg function (17). Furthermore, Bregs can influence disease progression and outcomes across various models, highlighting their critical role in immune regulation (17–20).

Distinct immune cell populations, including B cells, can also be differentiated by their metabolic profiles. Naïve B cells maintain low metabolic activity in both glycolysis and oxidative phosphorylation, but upon activation via antigen recognition or CD40/CD40L interaction, they undergo a significant metabolic shift, upregulating multiple pathways to support their functional requirements (1, 21). In contrast, Bregs exhibit lower glycolysis and oxidative phosphorylation rates but rely on higher fatty acid oxidation, a feature that may support their immunoregulatory function (22).

This study aimed to characterize the effects of ASNase treatment on B cell phenotype, antigen-presenting cell (APC) function, and metabolism. Additionally, we examined the ability of exogenous asparagine and glutamine supplementation to rescue ASNase-treated B cells, thus shedding light on the metabolic and functional consequences of ASNase exposure in non-malignant B cells.

## Material and Methods

### Sample preparation

Peripheral blood mononuclear cells (PBMCs) were isolated from healthy donor (HD) peripheral blood (PB) using density gradient centrifugation over Pancoll (Pan Biotech, Aidenbach, Germany). B cell enrichment was performed by positive immunomagnetic selection using CD19 microbeads (Miltenyi Biotec, Bergisch-Gladbach, Germany) according to the manufacturer’s protocol. T cells were purified by negative immunomagnetic selection using an EasySep T-cell enrichment kit (StemCell Technologies, Grenoble, France) according to the manufacturer’s protocol.

### Cell culture and drug treatment

PBMCs or B cells were cultivated at a cell density of 1-3x10^6^ cells/mL in 24 well plates in Iscovés modified Dulbecco medium (IMDM) (GIBCO by Life Technologies, California, United States) containing varying levels of glutamine and asparagine supplemented with 10% heat-inactivated human serum (obtained from donors), penicillin and streptomycin (GIBCO by Life Technologies, California, United States), 5 mg /mL insulin Actrapid (Novo Nordisk Pharma GmbH, Mainz, Germany), and 0.63 µg/mL Cyclosporin A (Sigma-Aldrich, Missouri, United States. The culture medium was additionally freshly supplemented with 50 U/mL recombinant human interleukin (IL)-4 (ImmunoTools, Friesoythe, Germany) and CD40 ligand multimer kit (Miltenyi Biotech, Bergisch-Gladbach, Germany) according to the manufactureŕs protocol. Cells were treated with 0-10 U/mL L-Asparaginase (Medac, Hamburg, Germany). All cells were cultivated at 37° C in a 5 % CO2 humidified atmosphere.

### Flow cytometry

Phenotype and proliferation were evaluated with flow cytometry using Gallios 10-colour flow cytometer (Beckman Coulter, California, United States) or Cytoflex LX flow cytometer (Beckman Coulter, California, United States). Used antibodies and dyes, see table **xx**.

For proliferation assessment cells were labelled with carboxyfluoresceinsuccinylimidyl (CFSE) or CellTrace violet (Thermo Fisher Scientific, Massachusetts, United States) according to manufactureŕs protocol.

For flow cytometric analysis 2x10^5^ cells were washed twice (550 rpm, 5 min) in Dulbeccós Phosphate Buffered Saline (DPBS) and staining was performed with 2 µg/mL of each antibody in 20 µL DPBS per sample. Samples were incubated for 20 minutes at 4° C in the dark, washed and resuspended in DPBS for sample evaluation.

### Microscopy

The microscopic evaluation of cluster formation was performed after 5 days of PBMC culture. Controls were taken on the first day of PBMC culture using the Leica Thunder imaging system (Leica Microsystems, Wetzlar, Germany).

### Mixed lymphocyte reaction

As stimulators, 2x10^4^ activated, irradiated B cells (26 Gy) were used for 10x10^4^ negatively selected, allogenic T cells in a final volume of 200 µL in 96-well round bottom plates. Prior to co-culture T cells were labelled with CFSE as previously described. Cells were harvested and after 5 days proliferation and phenotype were evaluated by flow cytometry.

### Enzyme-linked immunosorbent assay (ELISA)

IL-10 secretion by B cells was evaluated using ELISA. Supernatants from PBMC culture were taken on day 5 and analysed using ELISA MAX (Biolegend, California, United States) according to manufactureŕs protocol. Absorption was measured with the Cytation 1 (Biotek Instruments, Vermont, United States).

### Extracellular flux assay (Seahorse)

B cells were treated with asparaginase for 18h and 1.5x10^5^ cells were plated on a poly-D-lysine coated 96-well Seahorse utility plate (Agilent, California, United States). Metabolic function was analysed by measuring basal metabolism and metabolism after oligomycin, carbonyl cyanide-p-trifluoromethoxyphenylhydrazone (FCCP), and rotenone/antimycin A with 2-deoxyglucose injections, each with a final concentration of 1 µM.

### Metabolomics by LC/MS Analysis

Water-soluble metabolites were extracted from cell pellets with 500 μL ice-cold MeOH/H2O (80/20, v/v) containing 0.01 μM lamivudine and 1 µM each of D4-succinate, D5-glycine, D2- glucose and ^15^N-glutamate as external standards (Sigma-Aldrich, St. Louis, USA). After centrifugation of the resulting homogenates, supernatants were evaporated in a rotary evaporator (Savant, Thermo Fisher Scientific, Waltham, USA). Dry sample extracts were redissolved in 100 μL 5 mM NH4OAc in CH3CN/H2O (50/50, v/v). 15 μL supernatant was transferred to LC-vials. For LC-MS analysis 3 μL of each sample was applied to a XBridge Premier BEH Amide (2.5 μm particles, 100 × 2.1 mm) UPLC-column (Waters, Dublin, Ireland). Metabolites were separated with Solvent A, consisting of 5 mM NH4OAc in CH3CN/H2O (40/60, v/v) and solvent B consisting of 5 mM NH4OAc in CH3CN/H2O (95/5, v/v) at a flow rate of 200 μL/min at 45°C by LC using a DIONEX Ultimate 3000 UHPLC system (Thermo Fisher Scientific, Bremen, Germany). A linear gradient starting after 2 min with 100% solvent B decreasing to 0% solvent B within 23 min, followed by 17 min 0% solvent B and a linear increase to 100% solvent B in 1 min. Recalibration of the column was achieved by 7 min prerun with 100% solvent B before each injection. Ultrapure H2O was obtained from a Millipore water purification system (Milli-Q Merck Millipore, Darmstadt, Germany). HPLC-MS solvents, LC-MS NH4OAc, standards and reference compounds were purchased from Merck.

All MS-analyses were performed on a high-resolution Q Exactive mass spectrometer equipped with a HESI probe (Thermo Fisher Scientific, Bremen, Germany) in alternating positive- and negative full MS mode with a scan range of 69.0–1000 m/z at 70K resolution and the following ESI source parameters: sheath gas: 30, auxiliary gas: 1, sweep gas: 0, aux gas heater temperature: 120°C, spray voltage: 3 kV, capillary temperature: 320°C, S-lens RF level: 50. XIC generation and signal quantitation was performed using TraceFinder V 5.1 (Thermo Fisher Scientific, Bremen, Germany) integrating peaks which corresponded to the calculated monoisotopic metabolite masses (MIM +/− H^+^ ± 3 mMU).

### Data and statistical analysis

Flow cytometry data were analysed using Kaluza Software (Beckmann Coulter, version 1.1), FlowJo (BD Biosciences, version 10) or Cytolution (Cytolytics, Tübingen, Germany). Evaluation of microscopy pictures was performed using Leica Thunder imaging software (Leica Microsystems, version xx) and CellProfiler (CellProfiler, version 4). Statistical analysis was performed using GraphPad Prism (GraphPad, version 9.3). Three or more groups were analyzed by Kruskal-Wallis with Dunńs multiple comparison test. Three or more groups with more than one variable were analysed using two-way ANOVA with Tukey’s multiple comparison test. P values ≤ 0.05 were considered statistically significant.

## Results

### ASNase treatment reduces CD40-B cell numbers and clusters via proliferation inhibition but not increased cell death

To αinvestigate the overall effect of ASNase, we performed activation and proliferation experiments in CD40L-activated/stimulated PBMCs upon different ASNase concentrations. Here, we observed that increasing ASNase levels from 0 U/ml to 10 U/ml lead to less homotypic B cell clusters (**Fig. 1 A**). In addition, ASNase reduced the size and number of B cell clusters in a dose-dependent manner with a concentration of 10 U/ml reducing the area and the number of clusters compared to PBMCs cultured without ASNase (**Fig. 1 B+C**). To further validate these findings, we measured the expression of lymphocyte function-associated antigen 1 (LFA-1) as a key gene in lowering the threshold for B cell activation (REF). Flow cytometry-based analysis showed that B-cells had a lower expression of LFA-1 when exposed to higher levels of ASNase, LFA-1 expression decreased in B cells cultured with rising levels ASNase (**Fig. 1 D**).

**Fig. 1:**
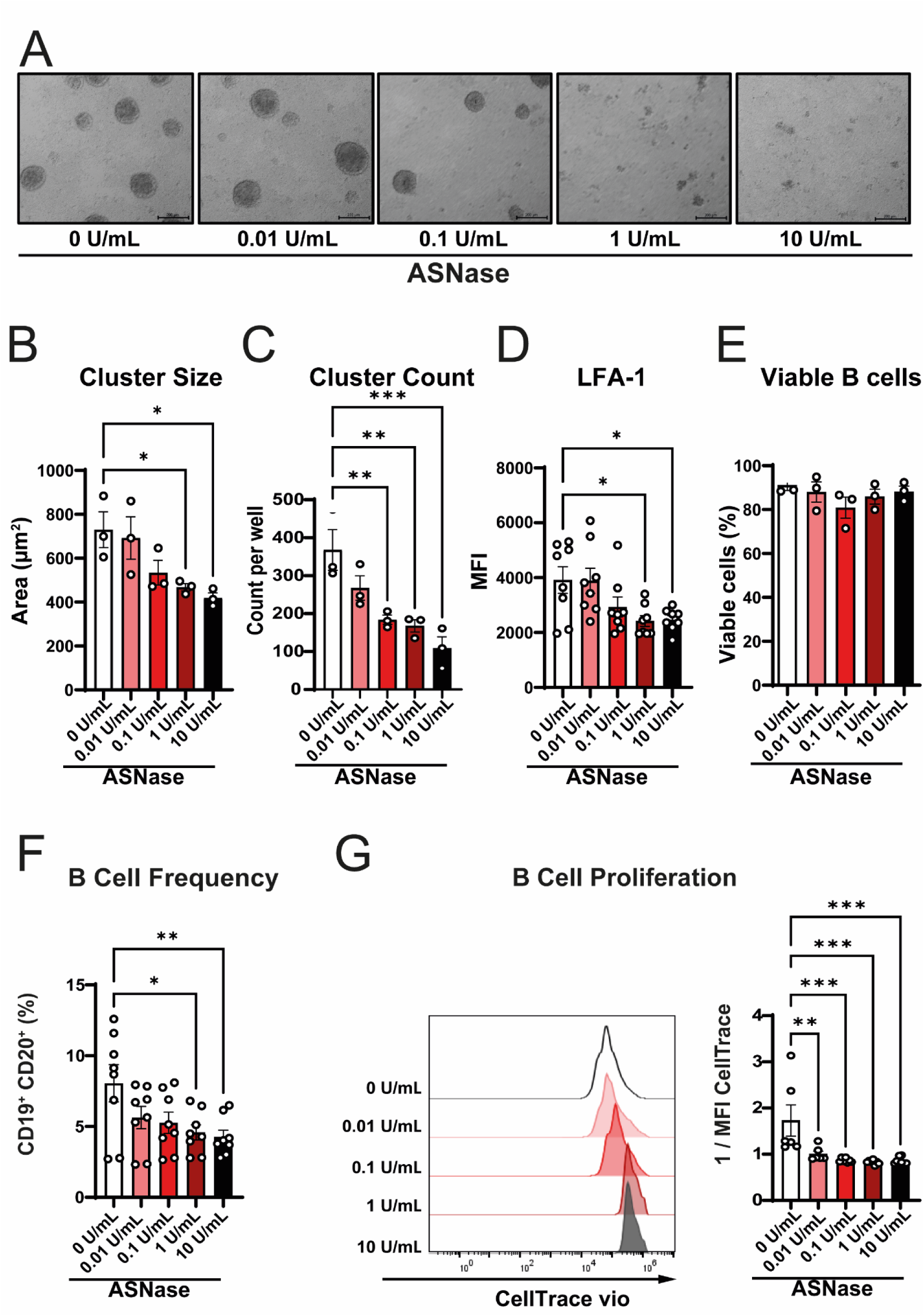
ASNase impairs B cell cluster formation and proliferation. Healthy donor PBMCs were cultivated for 5 days and activated with CD40L multimer. Cultivation included increasing ASNase concentrations. **A** Representative microscopic images of cluster formation. **B** Cluster sizes and **C** cluster counts of the respective images were analyzed using CellProfiler **D** LFA- 1 expression on CD19^+^ CD20^+^ B cells on day 5 of PBMC culture. **E** B cell frequency in PBMC culture on day 5 depending on ASNase concentration. **F** Representative histogram of CTV intensity and inverted MFIs showing B cell proliferation. All experiments were performed in biological triplicates or quadruplicates. Summary data is presented as mean with standard error of the mean. Statistical significance was calculated with one-way ANOVA (reference 0 U/ml).

While we did not observe any effects on B cell viability (**Fig. 1 E**) after an in-vitro culture time of 5 days, the addition of ASNase decreased B cell frequency (**Fig. 1 F**). Further proliferation assessments of staining revealed that treatment with ASNase stunted proliferation (**Fig. 1 G**). Here, culturing in lower ASNase concentrations, B cells partially proliferated, whereas a proliferation was nearly undetectable at higher ASNase concentrations.

Collectively, our results show a dose-dependent effect of ASNase on the activation and proliferative capacity on HD B cells without affecting viability.

### Antigen-presenting phenotype and function are reduced by ASNase treatmen

To assess if changes in the immune phenotype accompanied the changes in B cell proliferation, we measured the expression of CD86, HLA-DR, CD21 and CD80 on B cells after five days of incubation with CD40L and increasing ASNase concentrations. These markers allow for the evaluation of B cell activation and their APC function. B cell phenotype was analyzed after 5 days of CD40L activation combined with different ASNase concentrations.

Representative samples were concatenated, analysed for multiple markers and visualized using tSNE dimensionality reduction (**Fig. 2 A**). Regarding the clustering of samples, we observed a gradient from 0 U/mL to 10 U/mL. An analysis of individual markers revealed that CD86 expression was highest in cells from the untreated control group and signficiantly decreased in ASNase conetrations exceeding 0.1U/ml (**Fig. 2 B**). A similar trend was observed with HLA-DR expression (**Fig. 2 C**). While low levels of ASNase treatment did not significantly impact HLA-DR expression, treatment with 0.01 U/mL ASNase led to a notable decrease, with further reductions observed in a dose-dependent manner. Additionally, CD21 expression was inversely correlated with increasing ASNase concentrations (**Supp. 2 A**), whereas no significant changes were detected in CD80 levels (**Supp. 2 B**). Next, we investigated the effect of ASNase on the ability to present antigens by pre-incubating B cells with ASNase for five days, followed by irradiating and co-culturing with donor-matched T cells (**Fig. 3 D**). The activation of T cells was measured by the fraction of CD25^+^ CFSE^-^ T cells. Here, pre-incubation with ASNase led to an impairment of T cell activation and proliferation starting at a concentration of 0.01 U/ml ASNase. Together, these results suggest that a lack of asparagine interferes with CD40L-mediated B cell maturation and antigen presenting functions.

**Fig. 2:**
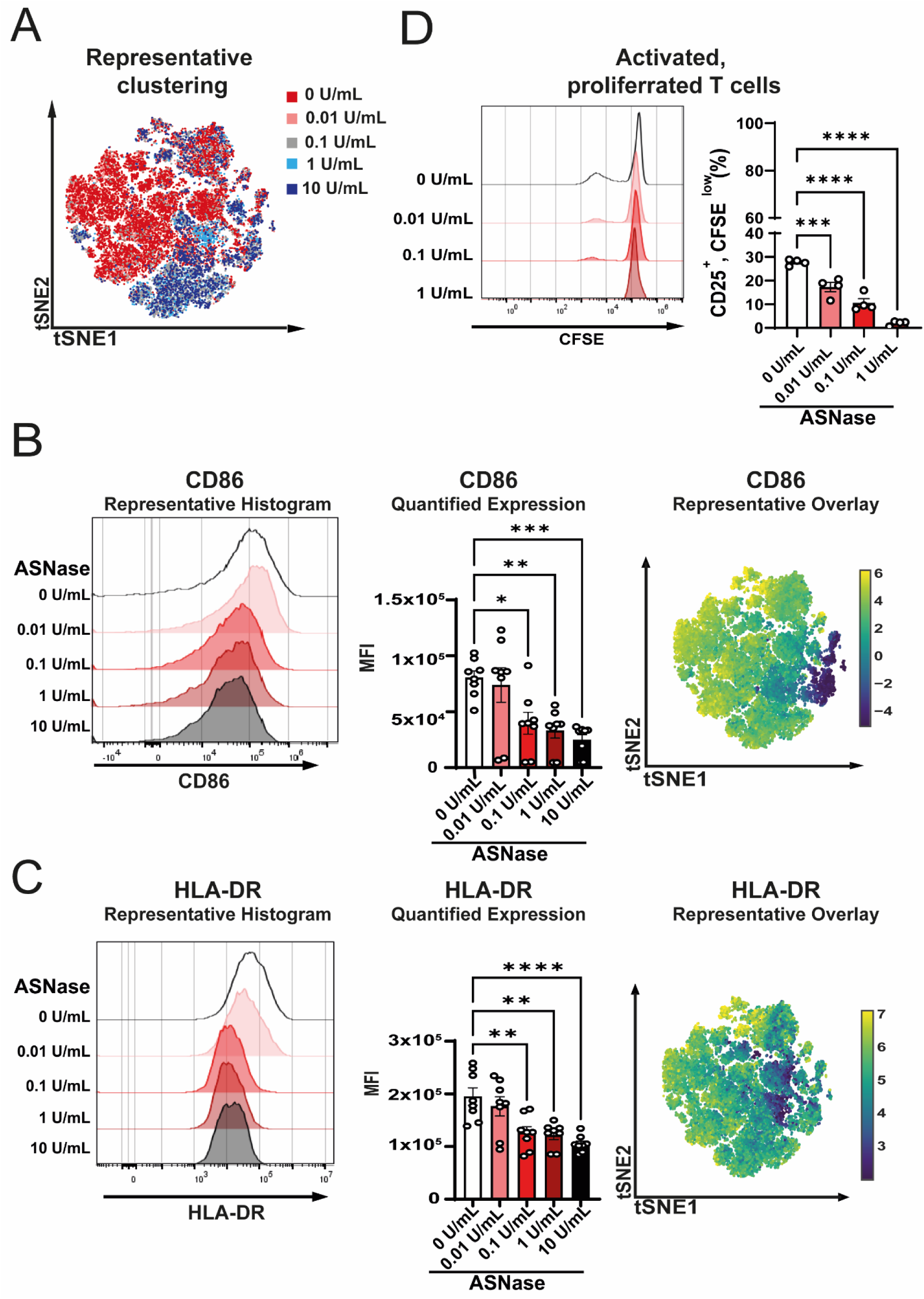
ASNase treatment hinders B cell activation and hinders APC function. Healthy donor PBMCs were cultivated for 5 days and activated with CD40L multimer. Cultivation included increasing ASNase concentrations. **A** Representative tSNE analysis of ASNase treated CD19^+^ CD20^+^ B cells. **B** Representative histogram of CD86 expression on CD19^+^ CD20^+^ B cells. Quantitative analysis of CD86 MFIs. Representative overlay of CD86 expression on treated CD19^+^ CD20^+^ B cells. **C** Representative histogram of HLA-DR expression on CD19^+^ CD20^+^ B cells. Quantitative analysis of HLA-DR MFIs. Representative overlay of HLA-DR expression on treated CD19^+^ CD20^+^ B cells. **D** Representative CFSE histogram of T cells co-cultivated with ASNase pre-treated B cells and quantification of activated, proliferated T cells. All experiments were performed in biological triplicates or quadruplicates. Summary data is presented as mean with standard error of the mean. Statistical significance was calculated with one-way ANOVA (reference 0 U/ml).

**Fig. 3:**
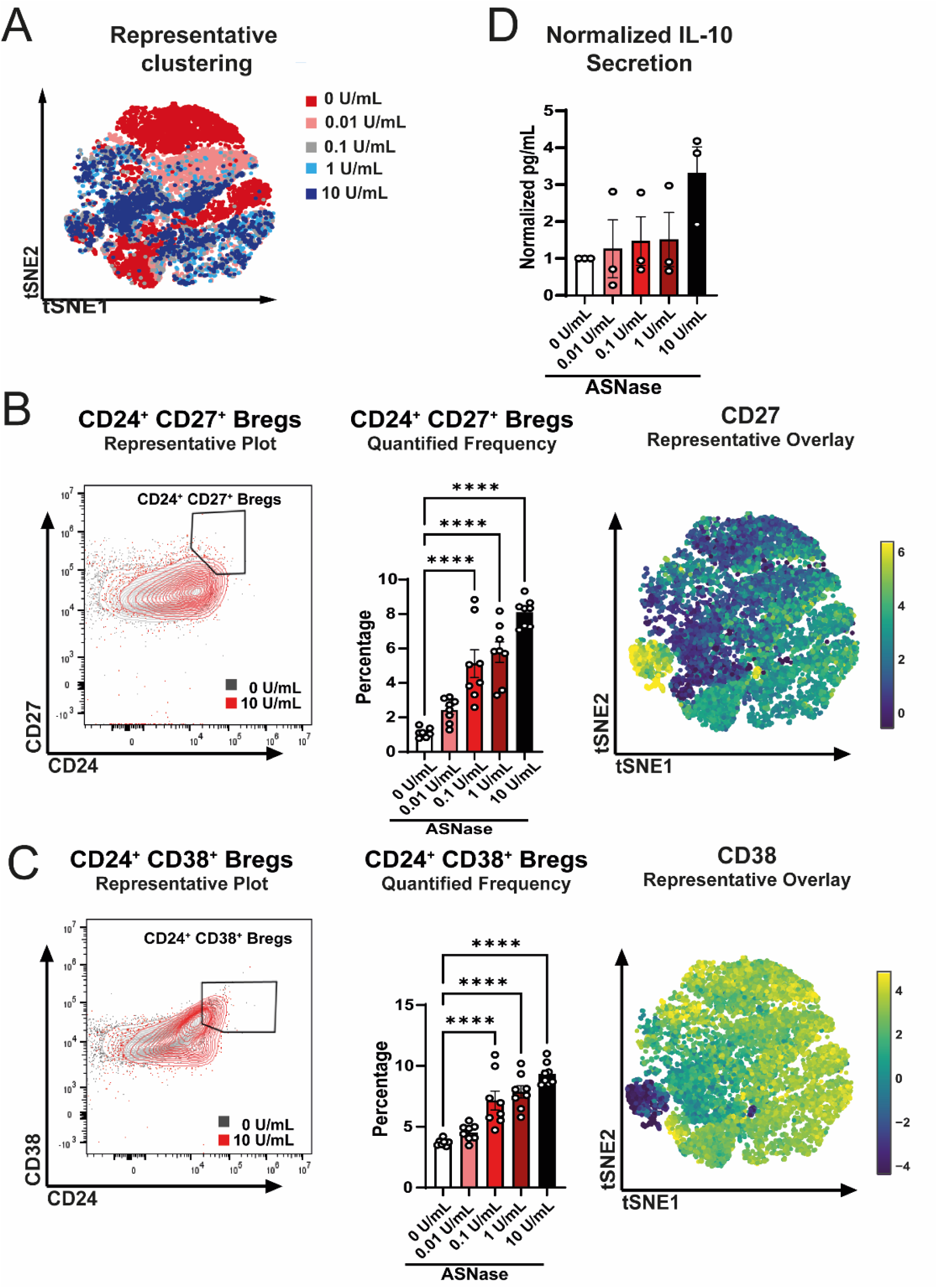
ASNase treatment induces Breg formation in a dose-dependent manner. Healthy donor PBMCs were cultivated for 5 days and activated with CD40L multimer. Cultivation included increasing ASNase concentrations. **A** Representative tSNE analysis of ASNase treated CD19^+^ CD20^+^ B cells. **B** Representative plots of CD24 and CD27 expression on CD19^+^ CD20^+^ B cells. Quantitative analysis of CD24^+^ and CD27^+^ Breg frequency. Representative overlay of CD27 expression on treated CD19^+^ CD20^+^ B cells. **C** Representative plots of CD24 and CD38 expression on CD19^+^ CD20^+^ B cells. Quantitative analysis of CD24^+^ and CD38^+^ Breg frequency. Representative overlay of CD38 expression on treated CD19^+^ CD20^+^ B cells **D** Normalized secretion of IL-10 into cell culture supernatant after days of CD40L activated B cell culture analyzed through ELISA. All experiments were performed in biological triplicates or quadruplicates. Summary data is presented as mean with standard error of the mean. Statistical significance was calculated with one-way ANOVA (reference 0 U/ml).

In the light of reduced APC function through ASNase treatment of B cells, we could not help but wonder, does ASNase promote the rise of Breg markers such as CD24, CD27, CD38, and TGF-β? Representative tSNE analysis revealed close clustering of non-treated and low-treated samples while rising ASNase concentrations shifted clusters (**Fig. 3 A**). CD24 and CD27, hallmarks for a Breg population when co-expressed, strongly increased with rising ASNase concentrations (**Fig. 3 B, S3 A**). Only a few Bregs are detected in the control samples while the Breg population strongly increases above 0.1 U/mL ASNase. The representative overlay underlines the findings. Further, the co-expression of CD24 and CD38 was analyzed, leading to similar results (**Fig. 3 C**). The co-expression rose after using 0.1U/mL ASNase levels. Besides, the secretion of IL-10 was analyzed from collected cell culture supernatants, further underlining the presence of a Breg phenotype when B cells are exposed to ASNase (**Fig. 3 B**). Normalized secretion shows a rise of measured IL-10 in a dose-dependent manner, Similarly, intracellular staining of TGF-β showed an increase in MFIs with rising ASNase concentrations (**Supp. 3 B**).

### ASNase induces metabolic reprogramming in CD40B-cells

Further analysis showed that granularity and size, both hallmarks of B cell activation, were strongly diminished by increasing ASNase treatment after reaching ASNase concentrations above 0.1 U/mL (**Fig 4 A, B**). To elucidate metabolic effects of ASNase treatment as a mechanism for the observed proliferation changes, we analyzed the mitochondrial and glycolytic metabolism by measuring the oxygen consumption and extracellular acidification in extracellular flux assays. Of note, untreated cells showed the highest oxygen consumption, while treatment with 0.1 U/ml and 1 U/ml ASNasegradually decreased oxygen consumption (**Fig. 4 C**). These differences were observed for both, the basal respiration (i.e., before the addition of FCCP) and the maximal respiration (i.e., after addition of FCCP), whereas the an ASNase concentration of 1 U/ml nearly halved oxygen consumption.

**Fig. 4:**
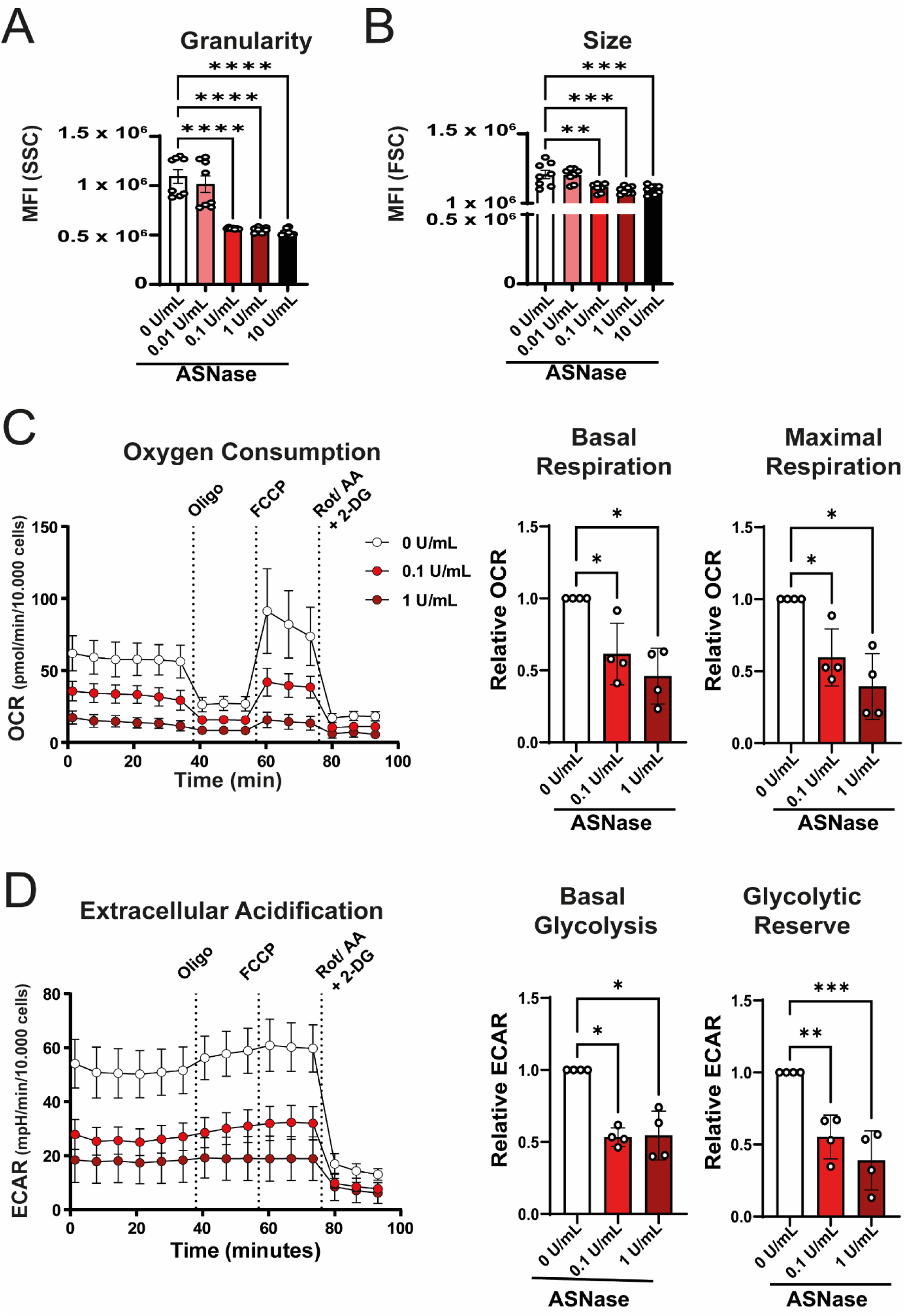
ASNase lowers metabolic function of B cells. Healthy donor PBMCs were cultivated for 5 days and activated with CD40L multimer. Cultivation included increasing ASNase concentrations. **A, B** Granularity and size were observed by flow cytometric analysis, bar graphs show the respective MFIs. **C, D** Purified B cells were metabolically analyzed using extracellular flux analysis after 18h ASNase treatment, following CD40L multimer expansion for 5 days. **C** Representative oxygen consumption rates of B cells following sequential oligomycin, FCCP, and rotenone + antimycin a + 2-DG treatment. Quantification of basal and maximal respiration. **D** Representative extracellular acidification rates of B cells following sequential oligomycin, FCCP, and rotenone + antimycin a + 2-DG treatment. Quantification of glycolysis and glycolytic reserve. All experiments were performed in biological triplicates or quadruplicates. Summary data is presented as mean with standard error of the mean. Statistical significance was calculated with one-way ANOVA (reference 0 U/ml).

We observed similar results for the glycolytic activity as measured by the extracellular acidification rate (**Fig. 4 D**). Here, basal glycolytic activity as well as the glycolytic reserve was impaired upon incubation with >0.1 U/ml ASNase. Moreover, we observed that the addition of oligmycin or FCCP did not alter ECAR in any of the conditions suggesting that CDD40L stimulation brought the B cells to their maximal glycolytic capacity. Glycolysis declined when cells were incubated with 0.1 U/mL or 1 U/mL ASNase. Similarly, glycolytic reserve shrunk through ASNase treatment. Oligomycin mediated ATPase inhibition and FCCP induced uncoupling of the electron transport chain lead to little change in ECAR as CD40L stimulated B cells are already at maximum capacity of their glycolytic rates.

We got further insights into the metabolic programs of ASNase-treated cells through metabolomic analysis. While treated cells contained significantly less L-asparagine the cellular content of the ASNase-cleaved product, L-aspartic acid was found in significantly higher concentrations. Similarly, the cellular levels of L-glutamine declined in a dose-dependent manner, whereas glutamic acid contents remained mostly unchanged (**Fig. 5 A**).

**Fig. 5:**
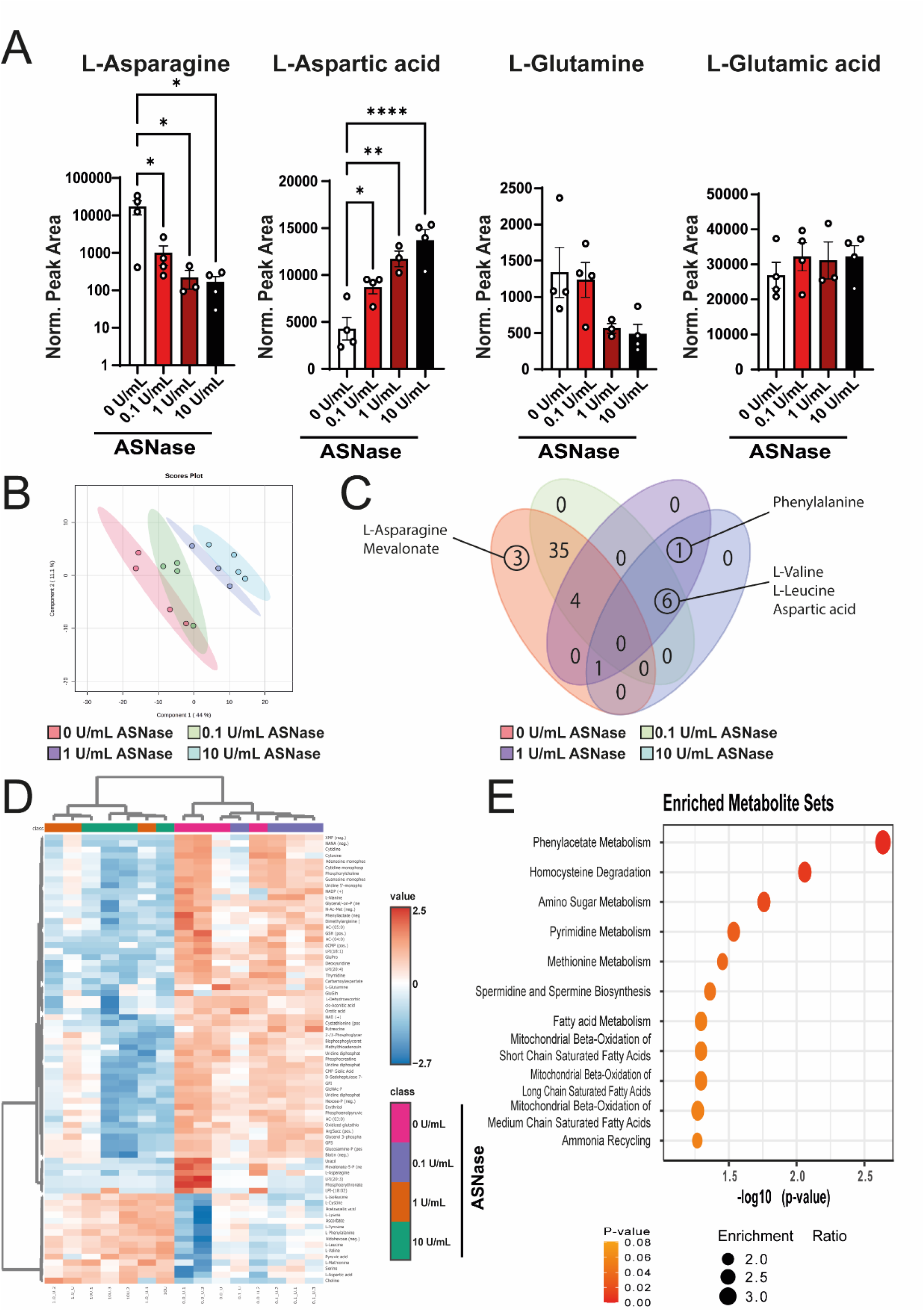
ASNase treatment alters metabolic profile of B cells. Healthy donor PBMCs were cultivated for 5 days and activated with CD40L multimer. Cultivation included increasing ASNase concentrations. **A** Cellular content of L-aspragine, L-aspartic acid, L-glutamine, L-glutamic acid as normalized peak areas. **B** PL-SDA plot of samples treated with different amounts of ASNase showing differential clustering. **C** Venn-diagram of upregulated metabolites. **D** Unsupervised clustering heatmap of all samples. **E** Enriched metabolites sets (0U/mL vs. 1U/mL). All experiments were performed in biological triplicates or quadruplicates. Summary data is presented as mean with standard error of the mean.

Together, these findings indicate that a decreased metabolic activity contributes to the detrimental effects of ASNase on B cell proliferation. Multivariate dimensionality reduction performed through partial least-squares discriminant analysis based on all measured metabolites revealed distinct, dose-dependent formation of clusters (**Fig. 5 B**). ASNase treatment induced a clear enrichment of certain metabolites in the differently treated samples as shown in **Fig. 5 C** depicting the amount of significantly enriched metabolites throughout the groups. L-Asparagine is enriched in untreated cells while treated cells took up significantly more other amino acids like valine, leucin, phenylalanine. Unsupervised clustering of singular samples reflects the distinct metabolic profiles further (**Fig. 5D**). Here, untreated and lowly treated samples (0.1 U/mL ASNase) cluster together whereas samples treated with higher concentrations (1-10 U/mL) share more of the same metabolic profile and therefore cluster together. Quantitative enrichment analysis comparing untreated B cells to B cells treated with 1 U/mL ASNase shows that untreated cells enrich for metabolite sets associated with proliferation and mitochondrial metabolism.

Our findings underline the mitigating effects of ASNase on B cell proliferation through lower cellular and metabolic activity due to lower rates in the main metabolic pathways.

### Exogenous amino acid supply rescues B cell cluster formation

Given the strong shifts in proliferation and APC function we asked if these effects are reversible through asparagine supply but also through glutamine supply as ASNase is not specific in cleaving only asparagine.

Similarly, to the previous experimental setting, B cells were now cultivated in medium containing an excess of either asparagine or glutamine or a combination of both. We observed the rescue of cluster size and formation, as shown in representative images (**Fig. 6 A**). Quantification showed that cultivation in IMDM alone containing physiological serum levels of asparagine and glutamine leads to a strong decrease in cluster size and formation (**Fig 6 B and C**). Recovery effects are observed through the exogenous supply of asparagine, whereas the stronger effect is seen with glutamine addition. The combination of both amino acids yields similar results. Accordingly, the size and granularity of B cells was analysed revealing the mitigating effects of exogenous amino acid supply. Size and granularity of B cells mostly closed in to non-treated cells when asparagine and glutamine were supplied.

**Fig. 6:**
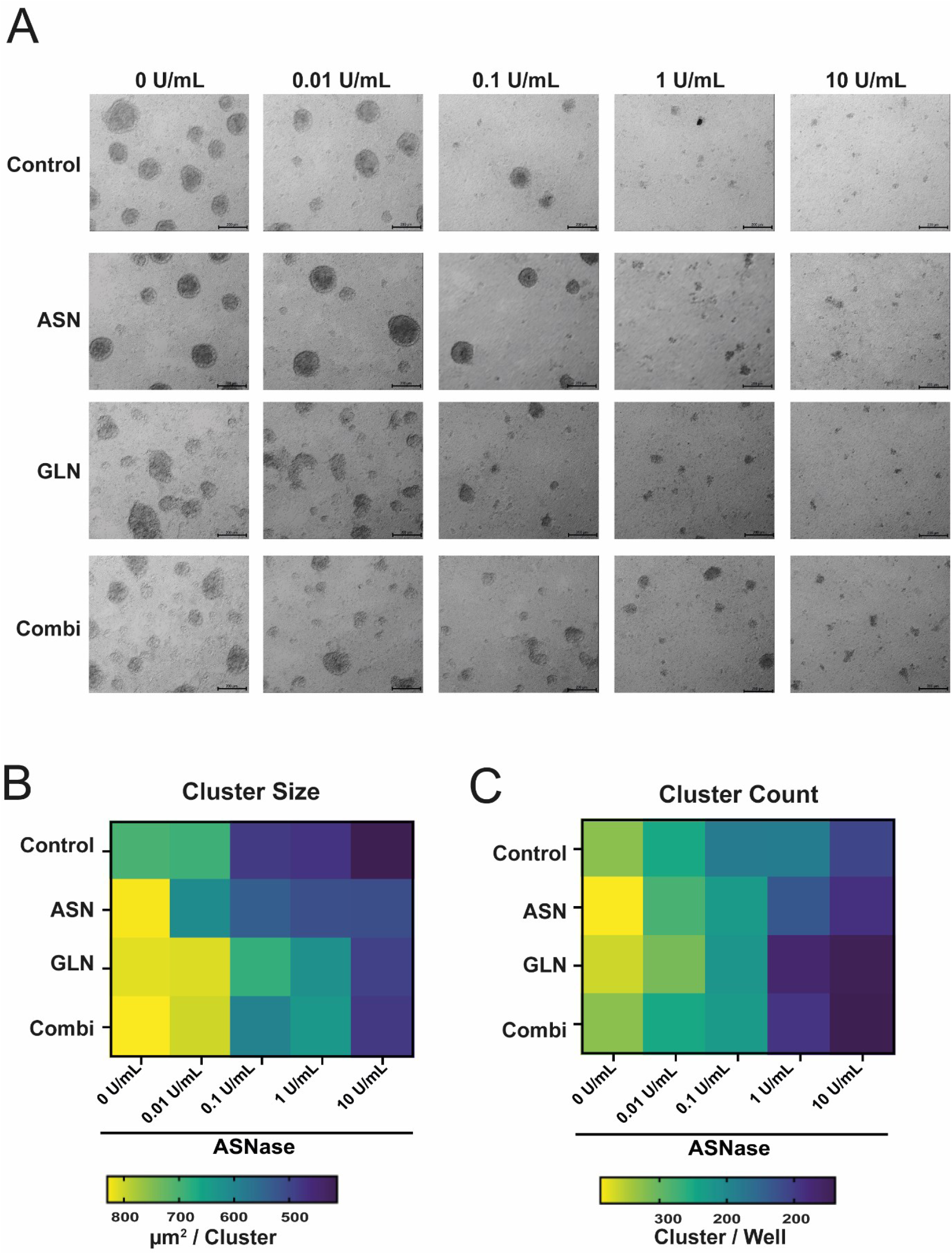
Exogenous supply of asn and / or glutamine rescues B cell cluster formation. Healthy donor PBMCs were cultivated for 5 days and activated with CD40L multimer. Cultivation included increasing ASNase concentrations. **A** Representative microscopic images of cluster formation in dependency of ASNase levels and exogenous supply of asparagine and glutamine. **B** Quantitative analysis of cluster size. **C** Quantitative analysis of cluster count. All experiments were performed in biological triplicates or quadruplicates. Summary data is presented as mean with standard error of the mean.

### Exogenous amino acid supply mitigates ASNase effect on B cell phenotype

In the same experimental setting the effects of exogenous amino acid supply were observed on phenotypic changes induced through ASNase treatment. The expression of CD86 was rescued through exogenous addition of asparagine and glutamine as well as through the combination of both (**Fig. 7A**). Through exogenous addition of asparagine and glutamine in low levels of ASNase (0.01 U/mL) expression of CD86 rose compared to non-supplemented medium. Supplementation of asparagine and/ or glutamine to cells treated with 0.1U/mL rescued CD86 expression to comparable levels as observed in untreated cells. Only in the non-supplemented medium CD86 is still decreased in the presence of 1 U/mL ASNase, while the addition of asparagine and/ or glutamine rescues the surface expression of CD86. No rescue at all can be observed when using 10U/ml (**Supp. 4A**). Similar effects can be observed with HLA-DR (**Fig. 7B**). In low levels of ASNase no significant difference in HLA-DR expression was seen between the normal IMDM and the supplemented medium, but also here a slight increase in HLA-DR expression can be observed. Incubation of cells in medium containing 0.1U/mL decreases HLA-DR levels in non-supplemented medium, supplementation reversed the effect. High levels of asparagine (1U/mL) decreased HLA-DR expression, yet supplementation of asparagine and combination of asparagine and glutamine reversed the impaired expression. Similarly, to CD86 no rescue function can be observed in 10U/mL of ASNase (**Supp. 4B**). We also investigated the effect of rescue mechanisms on CD24^+^ CD38^+^ Breg frequency (**Supp. 4C**). When looking at CD24^+^ CD38^+^ Breg frequency in low levels of ASNase glutamine and the combination of asparagine and glutamine even reduce CD24^+^ CD38^+^ Breg frequency when comparing to 0U/mL ASNase. While in 0.1U/mL ASNase alone does not show any effect on Breg frequency glutamine and the combination of asparagine and glutamine rescue Breg frequency to similar levels as in control. In higher doses (1U/mL) only glutamine addition rescues Breg frequency, asparagine alone and the combination of asparagine and glutamine fail to do so. Hence the B cell dependent T cell activation is also altered (**Fig. 7C**). In low levels of ASNase treatment (0.01U/mL) asparagine, glutamine, and the combination rescue APC function as observed through MLR experiments and the outcome of activated, proliferated T cells. In higher doses (1U/mL) the effect is only partially mitigated with the combination of asparagine and glutamine showing the highest efficacy.

**Fig. 7:**
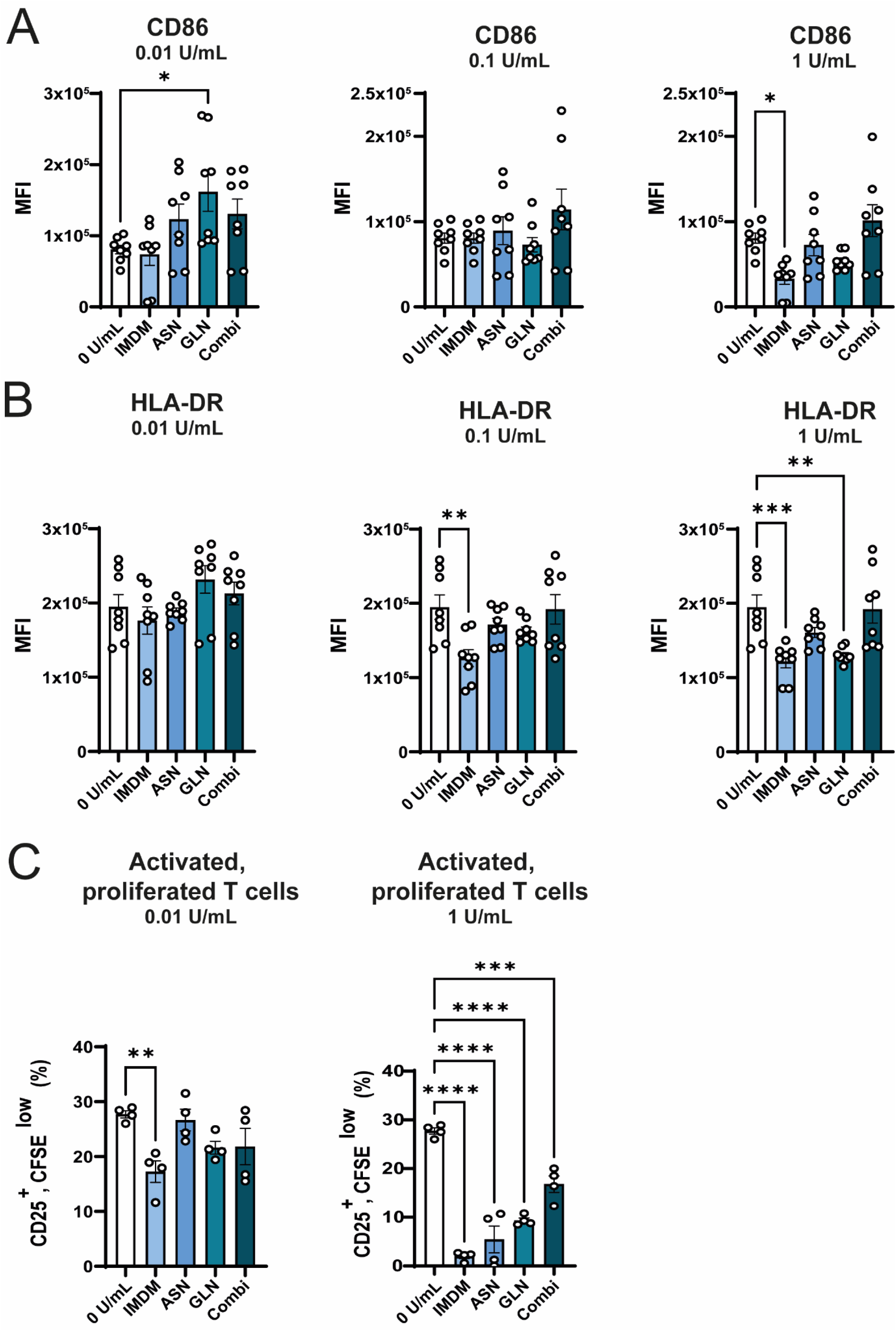
Exogenous supply of asparagine and / or glutamine rescues B cell APC function and counteracts Breg development. Healthy donor PBMCs were cultivated for 5 days and activated with CD40L multimer. Cultivation included increasing ASNase concentrations. **A** Quantitative analysis of CD86 MFI in different media under rising ASNase concentrations. **B** Quantitative analysis of HLA-DR MFI in different media under rising ASNase concentrations. **C** Quantification of activated, proliferated T cells activated in MLR through ASNase preincubated B cells in different media. All experiments were performed in biological triplicates or quadruplicates. Summary data is presented as mean with standard error of the mean. Statistical significance was calculated with one-way ANOVA (reference 0 U/ml).

### Metabolic programs mirror ASNase treatment and rescue through exogenous amino acid supply

While supplementation revived B cell clusters and phenotype we further aimed to analyze the underlying metabolic mechanism. The addition of asparagine or the combination with glutamine proved effective to bring up asparagine levels in the cells, further elevating aspartic acid acid levels as well, as competitive inhibition of ASNase led to a rise in the medium (**Fig. 8A**). Similarly, we observed addition of glutamine to elevate glutamine and glutamic acid levels in the cells. PLS-DA analysis shows a clear separation of different ASNase concentrations when medium is supplemented with asparagine, while the overlap grows through addition of glutamine (**Fig. 8B**). Quantification of significantly overrepresented metabolites where all treatment groups were compared with each other shows a separation of cells treated with low doses (0 – 0.1 U/mL) and samples treated with higher doses (1- 10 U/mL) ASNase. Pathway analysis conducted with these sets of metabolites show that samples treated with low doses enrich for pathways associated with proliferation while samples treated with higher doses of ASNase use recycling pathways like phenylalanine and tyrosine metabolism and ammonia recycling to compensate metabolic dysregulation.

**Fig. 8:**
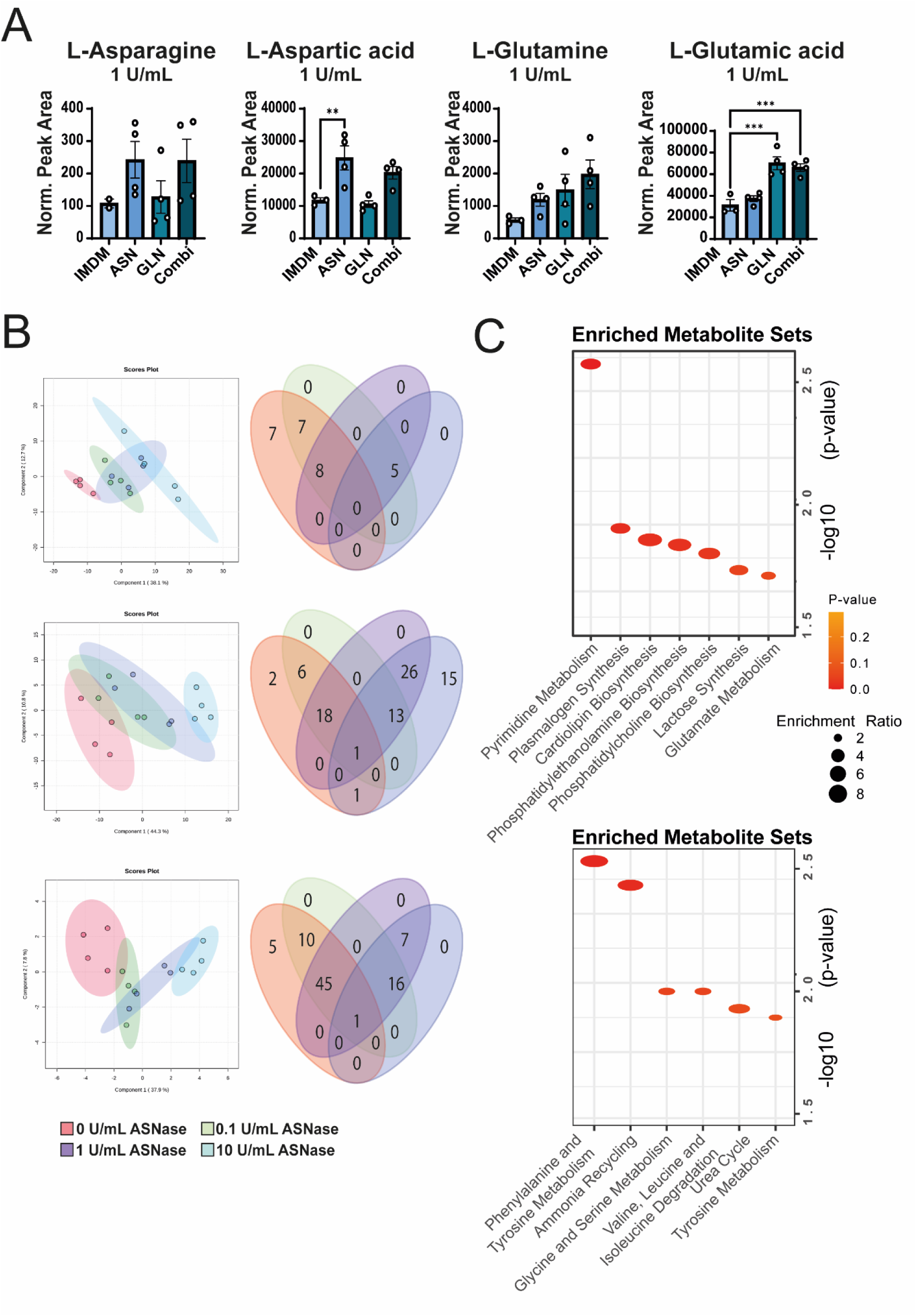
Exogenous supply of asparagine and / or readjusts metabolic profile. Healthy donor PBMCs were cultivated for 5 days and activated with CD40L multimer. Cultivation included increasing ASNase concentrations. **A** Normalized peak area of detected L- asparagine, L-aspartic acid, L-glutamine, L-glutamic acid detected in ASNase-treated cells (1 U/mL) and cultivated in supplemented media. **B** PL-SDA plots depicting samples treated different ASNase concentrations in the respective medium and venn-diagramm showing ANOVA testing results for observed metabolites. **C** Enriched metabolite sets based on overrepresented metabolites found in combination media (circled in **Fig. 8B**). All experiments were performed in biological triplicates or quadruplicates. Summary data is presented as mean with standard error of the mean. Statistical significance was calculated with one-way ANOVA (reference 0 U/ml).

## Discussion

Here in this study we investigated the effects of ASNase, a widely used therapeutic in B-ALL lowering asparagine and glutamine serum levels, on healthy B cells and showed adverse effects for B cell proliferation, APC function while observing increased regulatory functions. In the present study, we demonstrate novel aspects of using ASNase, focusing on the effects on healthy B cells. Our findings indicate that ASNase impairs B cell proliferation, skews B cells to a regulatory phenotype, and decreases antigen presentation. Of note, these effects seem to be reversible when glutamine and asparagine are supplemented in vitro. Thus, this study introduces new side effects of ASNase, extending known adverse effects like hepatotoxicity, fatigue, and pancreatitis, which unlike the other might be exploited clinically.

Healthy B cells possess functional asparagine synthetase and are seemingly not exclusively reliant on extracellular supply which concludes the low proliferation rather than a cytotoxic effect we observed on healthy B cells treated with ASNase. Most cellular mass during proliferation stems from amino acids (23), which we observed in treated B cells. Similarly to T cells, B cells upregulate their metabolism following activation (24). Unlike in B cells, as we have observed, in T cells, restriction of asparagine metabolism led to increased activation and increased anti-tumor immunity(25, 26). Whereas others observed the contrary (27, 28), even to the extent that ASNase treatment of T cells hindered metabolic reprogramming as it is observed in GC B cells (29, 30). As recent studies highlighted, asparagine proves crucial for B cell function (30). The compelling pre-print shows that highly active germinal center B cells rely in asparagine for homeostasis and function. Asparagine depletion, either through deletion of asparagine synthetase or environmental depletion of asparagine leads to decreased metabolic states resulting in poor exertion of GC function. Therefor adding to our observations that fully functional responses require that cellular metabolic needs are met accordingly. We showed that impaired asparagine uptake leading to decreased metabolic profiles comes with the decrease of APC function and increase of Breg phenotype. Classic APCs like dendritic cells and macrophages also rely on amino acid metabolism to exert APC function fully. Imbalanced plasma amino acid levels disturb fatty acid metabolism of dendritic cells observed in cirrhotic patients, causing delayed maturation and impaired APC function (31). Similarly to our findings in B cells, it was shown that ASNase disturbs macrophage function, dampening phagocytosis, proliferation, and antigen presentation (32). Hence, our findings in B cell APC function coincide with observations made in classical APCs. As exogenous asparagine supplementation hinders catalytic activity (33), thus overcoming the scarcity effects on B cells and their asparagine metabolism restoring healthy B cell function. As an unspecific target of ASNase activity glutamine levels are also decreased through ASNase treatment, which aids in treatment of ALL (34) but also impairs healthy B cell function (35). Therefore, reversibility of adverse effects is prominently mediated by glutamine supplementation and functions without asparagine supplementation as functional asparagine synthetase is present in healthy B cells catalyzing the amidotransferase reaction of aspartate and glutamine to asparagine(7) restoring regular B cell metabolism. We observed glutamine supplementation to be a major factor in reducing ASNase-mediating effects, similarly other studies observed that asparagine deficiency can not be compensated fully in glutamine scarce environments making asparagine conditionally essential (36). Here, healthy B cells with intact asparagine synthetase in combination with a glutamine supplemented environment can clearly compensate for the loss of asparagine.

Clinically, ASNase is used for B-ALL treatment only, due to its severe side effects ruling out its use when not absolutely necessary. Yet, we showed immunomodulatory effects already in low doses thus making ASNase a candidate drug for dampening autoimmunity. Impairment of CD24^+^, CD27^+^, CD38^+^ B cells shows increasingly adverse effects in autoimmunity in the context of multiple entities like Lupus, intestinal inflammation or rhinitis (37–39). (In the same context the question of increased infections arises, yet no data set is available showing correlation between ASNase treatment and rising infections due to its immunoregulatory features.Further, impairment of humoral immunity through ASNase was only described in mice (9, 11), data on human samples remain unavailable.) Conclusively, we showed the immunomodulatory effect of ASNase on B cells mediated by metabolic shifts in the cellular program. In the light of the results observed already through low-dose treatment we hypothesize a future exploration of ASNase as an immunomodulatory drug with manageable side-effects.

The study is limited by the defined levels of *in vitro* setting, we mimicked physiological glutamine and asparagine levels in the control medium but the variety of nutrients present makes it impossible to account for putative *in vivo* changes to other nutrients impacting cellular performance. Yet, this study adds valuable insights into B cell metabolism.

## Acknowledgements

We would like to thank Lucia Guldner for critical reading of the manuscript. We acknowledge the Core Facility Flow Cytometry at the Biomedical Center, Ludwig-Maximilians-Universität München, with Benjamin Tast and Can Pinar from Cytolution, Tübingen for assistance with data analysis and excellent technical support. This work was supported by the following grants and organizations: German Research Foundation, SFB-TRR338, DKTK, BZKF, Thomas Kirch Stiftung, EU Maria Skolodowskaya Research Network (T-OP),

## Author contributions

Conceptualization: AH, MG-M; Investigation: AH, MG-M, PB, NH, KF, DMCDS, KR, MF, AT, JT, AC, LV, SO; Formal Analysis and Visualization: AH, LV, MK; Methodology: AH, MG-M, ST; Writing Original Draft: AH, DMCDS; Writing Review and Editing: AH, DMCDS, ST, HS, MvB. All authors read and approved the final manuscript.

---

**Supplemental Fig. 1:**
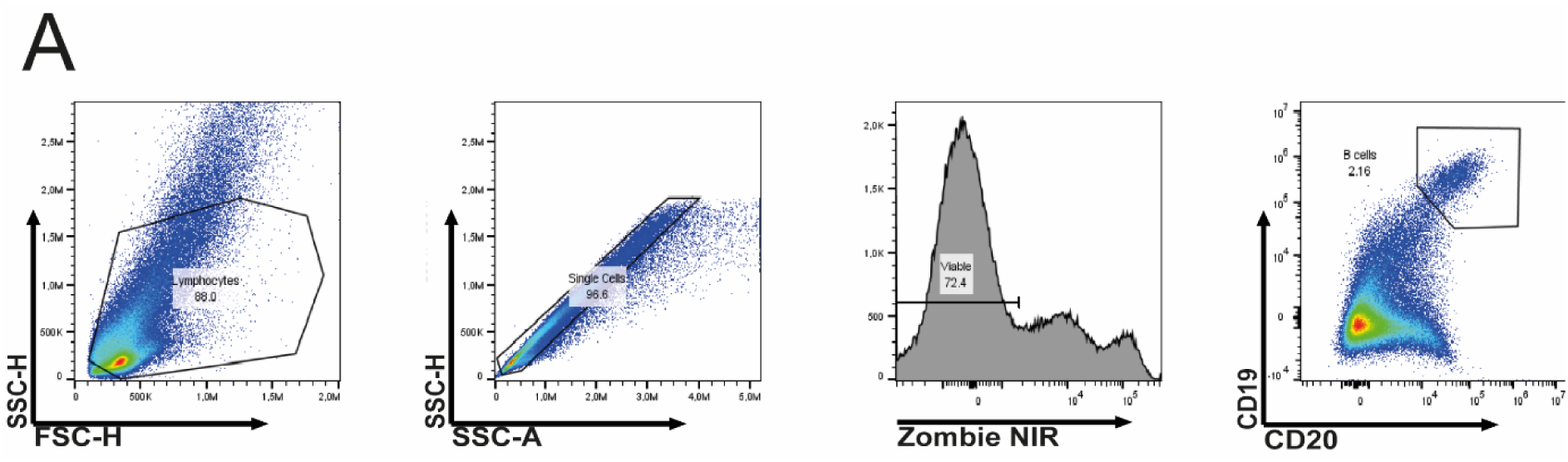
Gating strategy. Flow cytometry samples were gated on lymphocytes using forward and sideward scatter, then single cells using sideward scatter area against height, then on viable Zombie^-^ cells, then on CD19^+^ CD20^+^ B cells.

**Supplemental Fig. 2:**
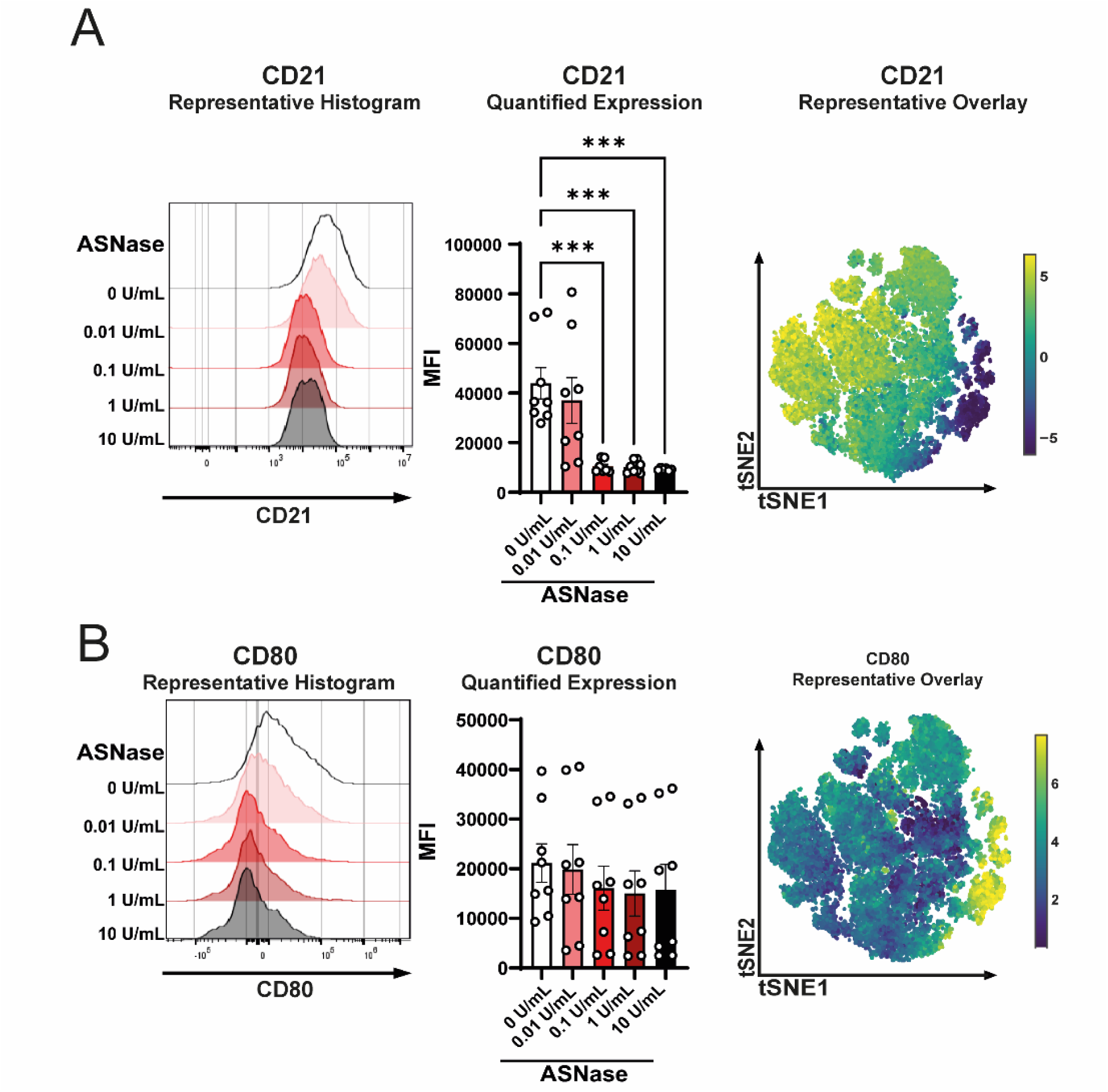
ASNase treatment hinders B cell activation and hinders APC function. Healthy donor PBMCs were cultivated for 5 days and activated with CD40L multimer. Cultivation included increasing ASNase concentrations. **A** Representative histogram of CD21 expression on CD19^+^ CD20^+^ B cells. Quantitative analysis of CD21 MFIs. Representative overlay of CD21 expression on treated CD19^+^ CD20^+^ B cells. **C** Representative histogram of CD80 expression on CD19^+^ CD20^+^ B cells. Quantitative analysis of CD80 MFIs. Representative overlay of CD80 expression on treated CD19^+^ CD20^+^ B cells. All experiments were performed in biological triplicates or quadruplicates. Summary data is presented as mean with standard error of the mean. Statistical significance was calculated with one-way ANOVA (reference 0 U/ml).

**Supplemental Fig. 3:**
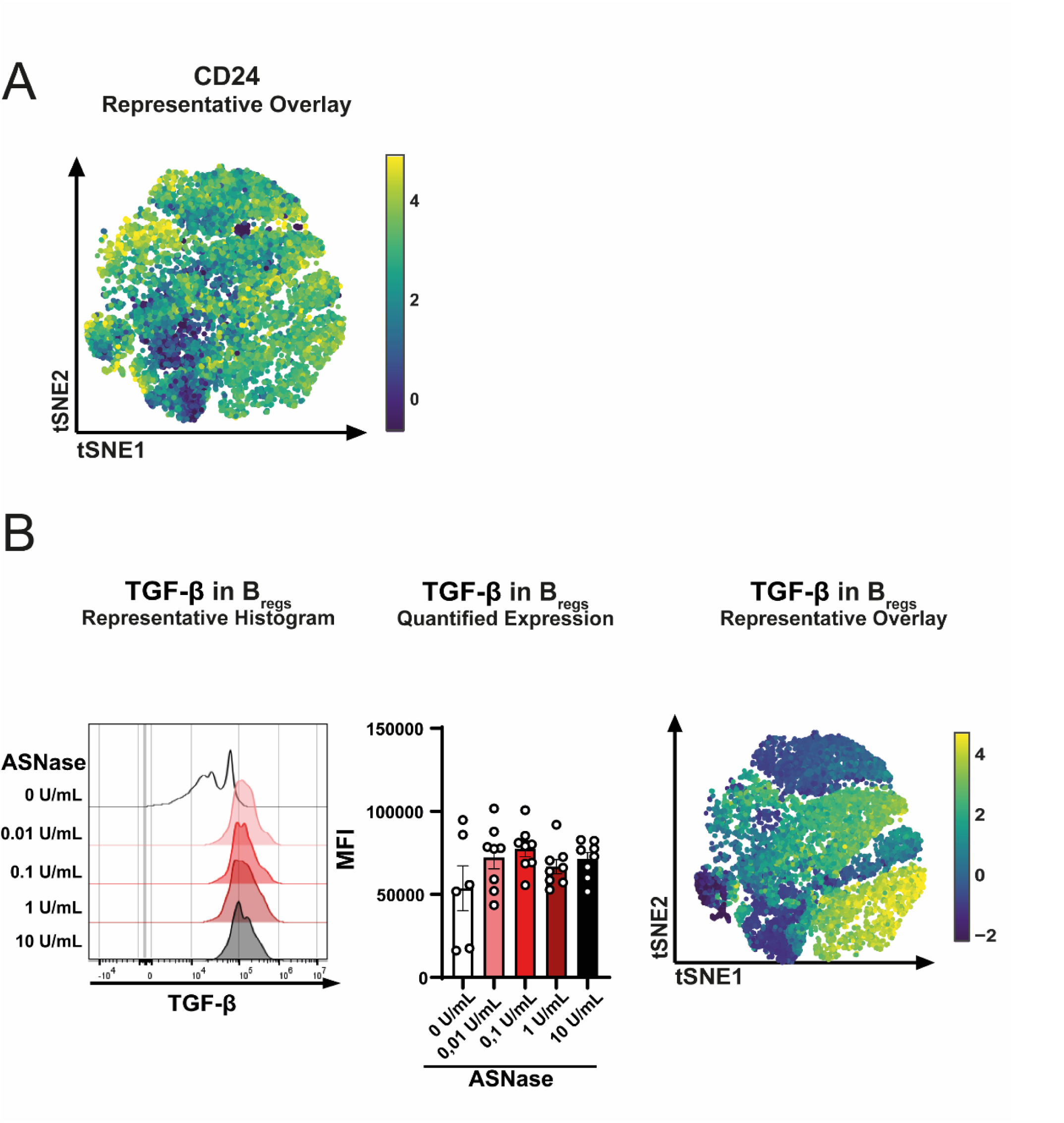
ASNase treatment induces Breg formation in a dose-dependent manner. Healthy donor PBMCs were cultivated for 5 days and activated with CD40L multimer. Cultivation included increasing ASNase concentrations. **A** Representative overlay of CD24 expression on treated CD19^+^ CD20^+^ B cells. **B** Representative histogram of TGFß expression on CD19^+^ CD20^+^ B cells. Quantitative analysis of TGF-ß MFI. Representative overlay of TGFß expression on treated CD19^+^ CD20^+^ B cells. All experiments were performed in biological triplicates or quadruplicates. Summary data is presented as mean with standard error of the mean. Statistical significance was calculated with one-way ANOVA (reference 0 U/ml).

**Supplemental Fig. 4:**
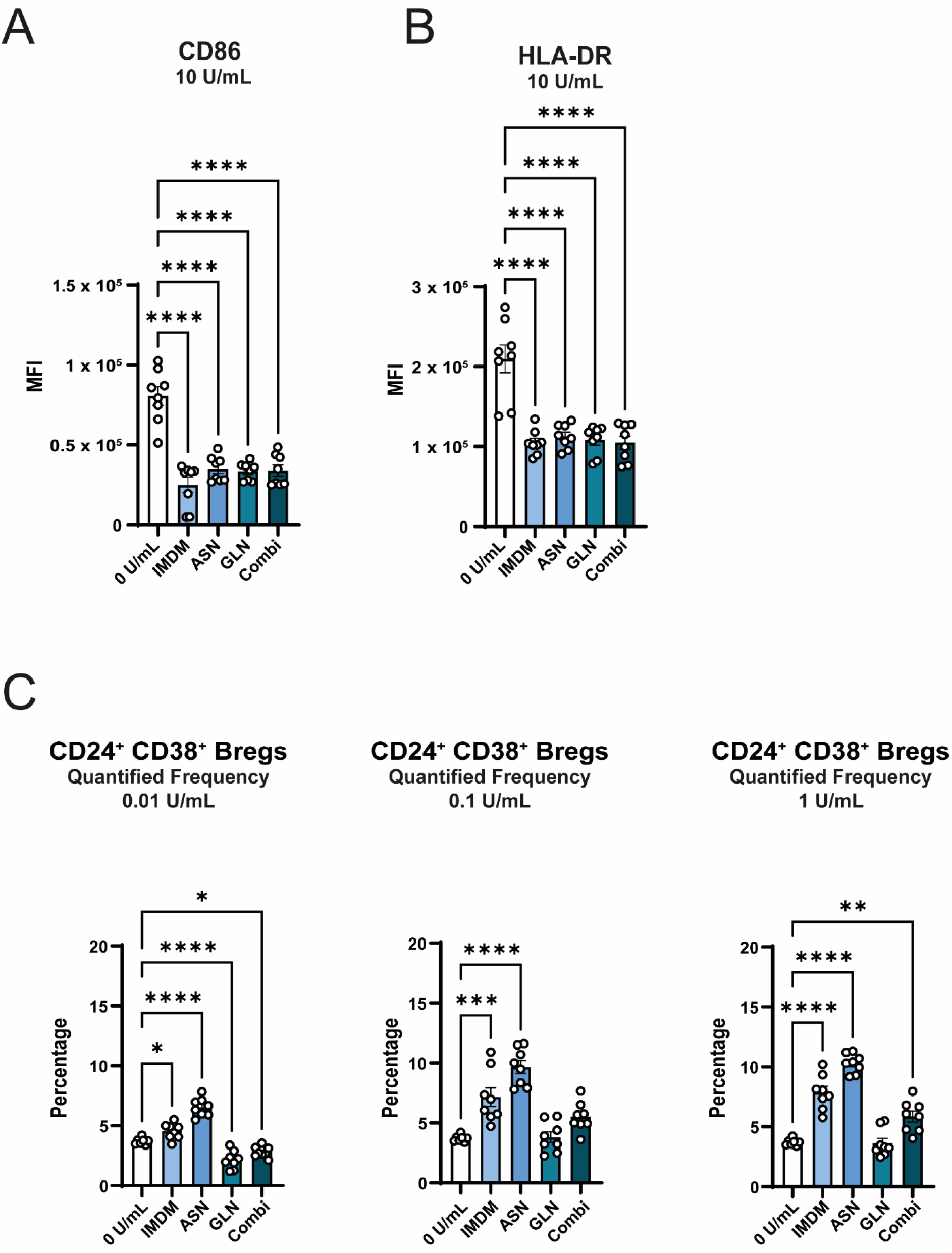
Exogenous supply of glutamine and / or glutamine rescues B cell APC function and counteracts Breg development. Healthy donor PBMCs were cultivated for 5 days and activated with CD40L multimer. Cultivation included increasing ASNase concentrations. **A** Quantitative analysis of CD86 MFI in different media under 10 U/mL ASNase. **B** Quantitative analysis of HLA-DR MFI in different media under 10 U/mL ASNase. **C** Quantification of CD24^+^ and CD38^+^ Bregs in different media under rising ASNase concentrations. All experiments were performed in biological triplicates or quadruplicates. Summary data is presented as mean with standard error of the mean. Statistical significance was calculated with one-way ANOVA (reference 0 U/ml).

